# COVID-19 as a continuous-time stochastic process

**DOI:** 10.1101/2023.03.08.531718

**Authors:** Irfan Lone, Pir Muzaffar Jan

## Abstract

In this article a mathematical treatment of Covid-19 as a stochastic process is discussed. The chance of extinction and the consequences of introducing new Covid-19 infectives into the population are evaluated by using certain approximate arguments. It is shown, in general terms, that the stochastic formulation of a recurrent epidemic like Covid-19 leads to the prediction of a permanent succession of undamped outbreaks of disease. It is also shown that one is able to derive certain useful conclusions about Covid-19 without consideration of immune individuals in a population.

## I. INTRODUCTION

The coronavirus disease 2019, also called Covid-19, is a fatal pandemic disease that started spreading around the world since the beginning of 2020 [1–5]. It is caused by a novel Severe Acute Respiratory Syndrome Coronavirus named SARS-Cov-2 [3, 6, 7]. The disease is believed to have originated from the Chinese city of Wuhan, perhaps after initially spreading into a closed community from a single infected individual, and has posed an extraordinary threat to global public health by infecting and killing millions worldwide [1–3]. Dynamic modeling is one of the time-tested mathematical tools utilized in the study of epidemics and both deterministic as well as stochastic approaches are in existence [8]. However, the transmission of disease between individuals in real world is inevitably a random process [9]. In fact, if the size of outbreak is significantly smaller than that of the total population a stochastic approach is a better choice for making useful predictions [9]. An understanding of Covid-19 dynamics through stochastic modeling should thus greatly assist in the control and prevention of this infectious disease [10]. Surprisingly, however, the stochastic modeling for Covid-19 has yet been relatively rare compared to its deterministic counterparts [9–16]. Probably the unique features of the outbreak are limiting the applicability of all existing models. For one, the determination of an accurate reproduction number for Covid-19 can be a challenging task as many asymptomatic infections may not get accurately accounted for [17]. Secondly, SARS-CoV-2 is more transmissible than SARS-CoV and MERS-CoV and the animal origin and cross-species infection route of SARS-CoV-2 are yet to be fully uncovered [3]. Finally, the large uncertainties arising from both poor data quality and inadequate estimations of model parameters (incubation, infection, and recovery rates) can propagate to long-term extrapolations of infection counts [18]. This unique transmission dynamics of Covid-19 has necessitated the development of novel stochastic models. On the basis of recent experimental studies of Covid-19, our treatment assumes that for a certain time period the disease infected individuals are asymptomatic and are not immediately infectious to the healthy persons. After a certain lapse of time, they turn into carriers and become infectious without showing any symptoms. These carrier individuals, after a certain time period, start showing symptoms of the disease and get isolated. They are either then cured and turn immune or else they die. Thus, our treatment is closely related to the so-called Susceptible–Exposed-Infected–Removed (SEIR) stochastic models [8]. For the sake of convenience of mathematical treatment, we make certain simplifying assumptions. First of all, we assume that latent period for Covid-19 is zero. This means that the infected individual becomes infectious to others immediately after the receipt of the infection. It is also highly convenient to assume that the length of the infectious period has a negative exponential distribution. Another quite reasonable assumption is that the chance of any susceptible becoming infected in a short interval of time is jointly proportional to the number of infectives in circulation and the length of the interval. This, in other words, means that the chance of one new infection in the whole group in a short interval of time will be proportional to the product of the number of infectives and the number of susceptibles, as well as the length of the interval. The transition probability will thus be a non-linear function of group size.

## II. BASIC THEORY CHARACTERIZING A STOCHASTIC EPIDEMIC

In our stochastic treatment, we represent the number of susceptibles that are still uninfected at time *t* by the random variable *R*(*t*) while as the number of infectives that are in circulation is represented by the random variable *S*(*t*). In this way, for a group of given total size, the number of removed cases gets fixed once the values of *R* and *S* are known and the process therefore becomes two-dimensional and is treated as follows. We denote the probability of having *u* susceptibles and *v* infectives at time *t* by *p*_*uv*_(*t*), i.e.

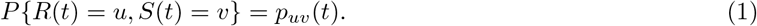

For the sake of convenience of mathematical treatment, we shall work with the following probability-generating function

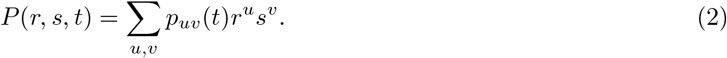

Let the joint probability distribution of relevant transitions in the interval Δ*t* be given by

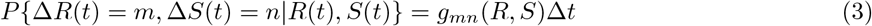

This excludes the case of both *m* and *n* jointly being zero. Now there are two types of possible transitions in the process. The probability of occurrence of a new infection in time Δ*t* is *αRS*Δ*t* and *u* decreases by one unit while *v* increases by one unit. The probability of removal is *βS*Δ*t* and *v* decreases by one unit. These transitions are represented in our treatment by (*m*.*n*) = (−1, +1) and (0, −1). The function *g*_*mn*_(*R, S*) then takes only the following two non-zero values:

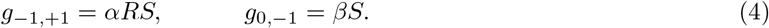

The necessary partial differential equation for *P*(*r, s, t*) is then given by

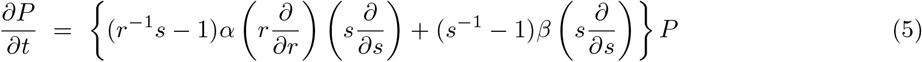

More succinctly,

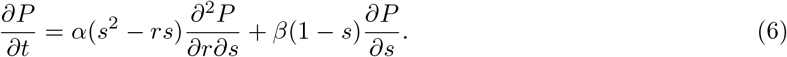

Changing the time scale to *τ* = *αt*, and writing 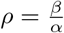 above equation becomes

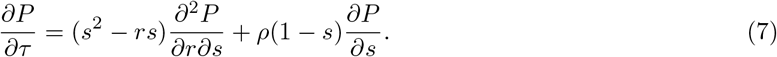

Let the process start at *τ* = 0 with *b* susceptibles and *a* infectives. The initial condition is then

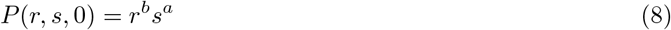

A direct solution of Eq. (7) is intractable but if we pick out the coefficients on *r*^*u*^*s*^*v*^ on both sides, or alternatively express *p*_*uv*_(*t* + Δ*t*) in terms of the probabilities at time *t*, we obtain the following set of differential equations

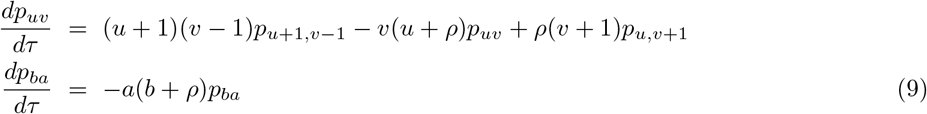

where 0 ≤ *u* + *v* ≤ *b* + *a*, 0 ≤ *u* ≤ *b*, and 0 ≤ *v* ≤ *b* + *a* with

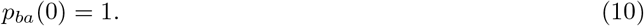

where it is assumed that any *p*_*uv*_, whose suffices fall outside the permitted range, is zero. The easiest way to handle the system of equations given by (9) is to make use of the Laplace transform [23] and its inverse, defined for any function *ϕ*(*t*) by the following relations

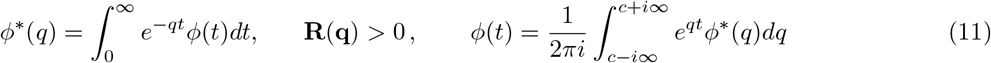

where *c* is positive and greater than the real parts of all singularities of *ϕ*\*(*q*). Suppose we use the Laplace transform in (11), writing *q*_*uv*_(*q*) for the transform of *p*_*uv*_(*τ*), i.e.

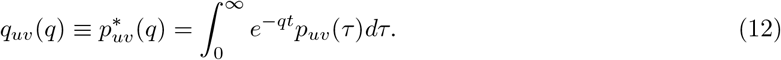

equations (9) can then be transformed into

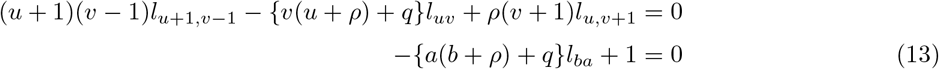

with the same ranges as before for *u* and *v*. The above treatment easily carries over into the case of Covid-19 as a recurrent pandemic. To this end we consider a group of susceptible individuals all mixing homogenously together. One or more of the individuals then contract Covid-19 which in due course is passed on to the other susceptibles. After the receipt of infectious material, a coronavirus, there is a latent period during which the disease develops purely internally within the infected person. The latent period is followed by an infectious period during which the infected person (or infective as he is then called) is able to discharge the virus in some way and probably communicate the disease to other susceptibles. Sooner or later actual symptoms of Covid-19 appear in the infective who is then removed from circulation among the susceptibles until he either recovers or dies. The removal of the infective brings the infectious period effectively to an end at least as far as the possibility of spreading the disease further is concerned. The time interval between the receipt of infection and the appearance of symptoms is called the incubation period. Covid-19 has an incubation period of 1–14 days, typically around 5 days, depending on the viral load [3].

## III. THE CASE OF COVID-19

A characteristic feature of Covid-19 is that each outbreak has a kind of epidemic behavior discussed above but in addition these outbreaks tend to recur with a certain regularity. Covid-19 is then in a sense endemic as well as epidemic. In order to construct a stochastic model of this recurrent behavior we start with the general stochastic epidemic of the previous section and modify it by having new susceptibles introduced into the population according to a Poisson process with parameter *σ*. This means that there are now three possible types of transition in time Δ*t*. As before, we use *R*(*t*) and *S*(*t*) to represent the number of susceptibles and infectives at time *t*, with probability *p*_*uv*_(*t*) given by equation (1). A new infection will occur in Δ*t* with probability *αRS*Δ*t*, and *u* decreases by one unit and *v* increases by one unit. A removal will occur with probability *βS*Δ*t*, and *v* then decreases by one unit. In addition, we may have the introduction of a new susceptible, with probability *σ*Δ*t*, and *u* will increase by one unit. In the notation of previous section, the transitions are represented by (*m, n*) = (−1, +1), (0, −1), and (+1, 0); and *g* = *αRS, g*_0,−1_ = *S* and *g*_1,0_ = *σ*. The required partial differential equation for *P*(*r, s, t*) is then given by

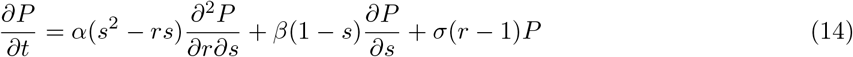

with the initial condition

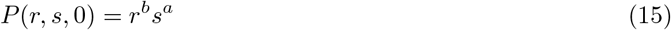

Just like Eq. (7), this basic equation for the process is intractable. One can use it to write down the differential-difference equations, like equations (9), for the individual probabilities *p*_*uv*_ but these too don’t yield to any investigation. In spite of these difficulties, certain fundamental properties of the process can be elucidated, at least approximately. It turns out that the threshold behavior of a general epidemic in a large group could be studied to some extent by observing that if *b* were large then at least initially the group of infectives was subject to a birth-and-death process with birth- and death-rates *αb* and *β*, respectively; and starting at *t* = 0 with *a* individuals. Suppose we ignore the loss of susceptibles due to infection, but take the arrival of new susceptibles into account by assuming these to follow a deterministic process with arrival-rate *σ*. The approximate birth- and death-rates for the group of infectives will then be *α*(*b* + *σt*) and *β*. We then have a birth-and-death process which is non-homogenous in time and for which we may take the birth- and death-rates *π*(*t*) and *σ*(*t*) to be

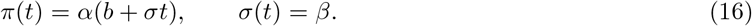

A general treatment of this process is a little involved. We can, however, easily handle the important aspect of extinction by first briefly investigating a simple kind of non-homogenous birth-and-death process as follows. Once again, we assume that only two types of transitions are possible, namely that the chance of any individual giving birth in time *t* is *π*(*t*)Δ*t*, and the chance of dying in Δ*t* is *σ*(*t*)Δ*t*. We thus have the transitions given by *m* = +1, −1; with *g*_+1_(*R*) = *π*(*t*)*S* and *g*_−1_(*R*) = *σ*(*t*)*R*. The required partial differential equation is then

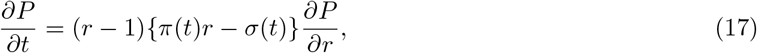

with initial condition

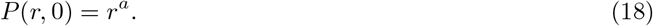

Equation (17) can be easily solved through a standard technique. We first write down the following subsidiary equations

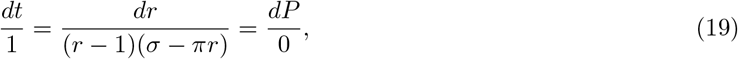

remembering that now both *π* and *σ* are functions of time, *t*. The first and third items in (28) immediately give one integral

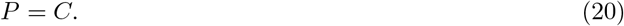

where *C* is an integration constant. Another integral can be found from the solution of

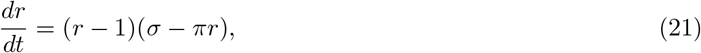

which simplifies somewhat by use of the following substitution

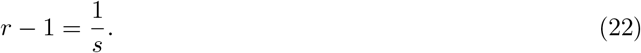

we find

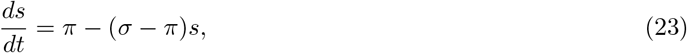

which now involves only a linear function of *s* on the right. The usual integrating factor for such an equation is *e*^*ρ*(*t*)^, where

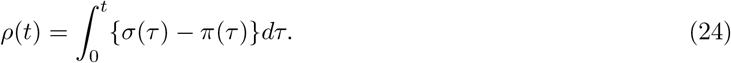

We can thus write

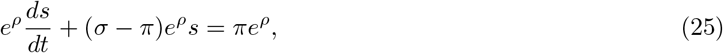

which, on integration with respect to *t*, yields

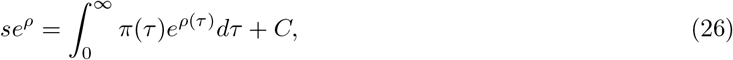

Alternatively,

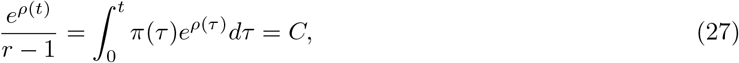

which is the required second integral. The general solution of (17) can therefore be expressed in the form

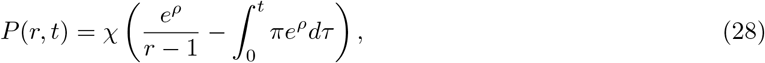

where the arguments of *π* and *ρ* have been suppressed for convenience. Putting *t* = 0 in (28), and using the initial condition (18), gives

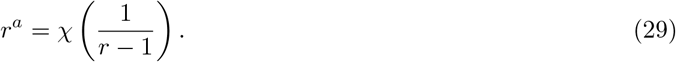

Writing *u* = (*r* − 1)^−1^ or *r* = 1 + *u*^−1^, gives the desired form of *χ* as

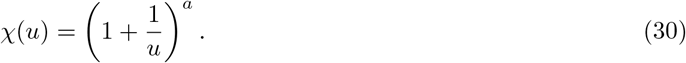

The final expression for the probability-generating function is therefore given by applying (30) to (28), i.e.

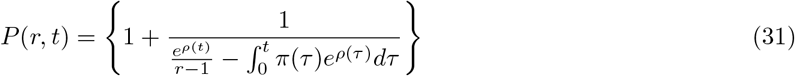

where *ρ*(*t*) is defined by (24). There is little difficulty in expanding (31) in powers of *r* to obtain the individual state probabilities. The general formula is a little complicated, but can be written as

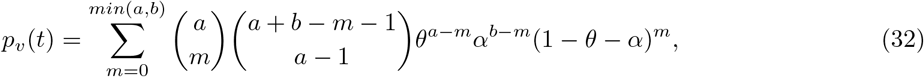

with

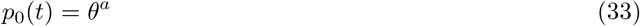

where

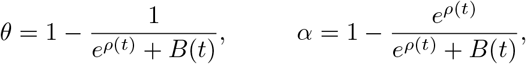

and

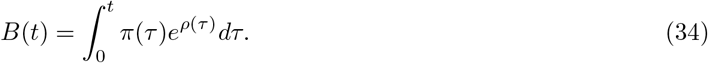

Now the chance of the process becoming extinct by time *t* is given by *p*_0_(*t*). From (35) this is

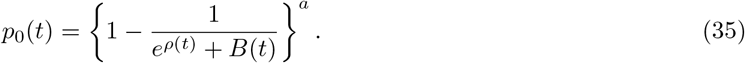

Now

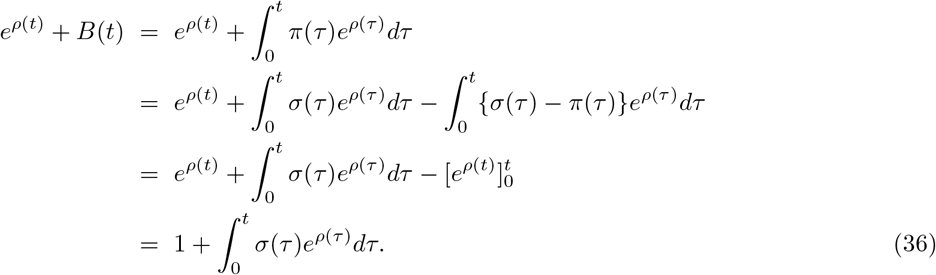

using (24). Substituting this in (35) gives

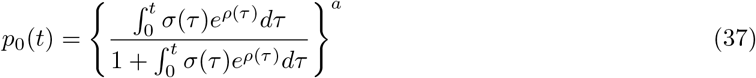

It is evident from (37) that the chance of extinction tends to unity as *t* → ∞, if and only if

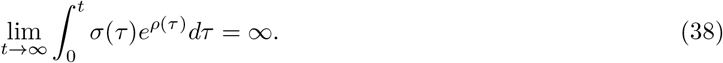

## IV. THE CHANCE OF EXTINCTION OF COVID-19

We can now calculate the chance of extinction of a recurrent epidemic like Covid-19 by making use of equations (16) and (24) to write

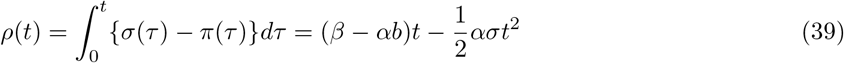

Thus, the chance of extinction *p*_0_(*t*), given by (37), can be written as

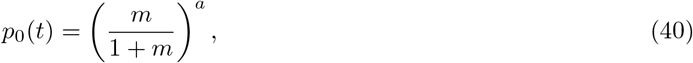

with

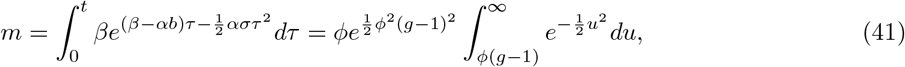

where

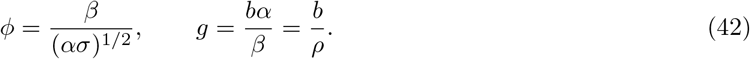

When *σ* = 0, the stochastic process reaches its threshold with a discontinuity in the chance of extinction at *ρ* = *b*. But if *σ* ≠ 0, there is no sharp cut-off at *ρ* = *b*. The probability of extinction will vary continuously, having the value

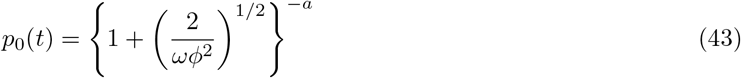

at *ρ* = *b*. If *ρ* is much larger than *b*, we shall have *g* → 0. If *ϕ* is large, *m* will be large, and *P* → 1. The consequences of introducing new Covid-19 infectives into the group, in order to trigger off a fresh epidemic as soon as the group of susceptibles has increased sufficiently, can be investigated to some extent by further approximate arguments. Thus, suppose the new infectives arrive at random with rate *ϵ*, so that the chance of a new arrival in Δ*t* is *ϵ*Δ*t*. Suppose also that at time *t* = 0 the number of susceptibles is negligible, but that it increases at a deterministic rate *σ*, which is much larger than *ϵ*. We neglect, as before, the ultimate reduction in the number of susceptibles when the disease begins to spread. Under these assumptions the effects of each newly introduced case of disease are independent of one another. Consider now the interval (*u, u* + Δ*u*). The chance of no new infective appearing is 1 − *ϵ*Δ*u*. The chance that a new infective will arrive is *ϵ*Δ*u*, when there will be a total of *σu* susceptibles. Any outbreak of disease resulting from this one case will be extinguished with probability *P*(*u*) given by

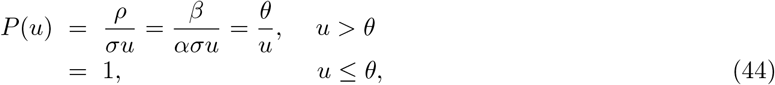

where *θ* is defined as *β/ασ*. Thus, the chance that no epidemic builds up from events occurring in Δ*u* is 1 − *ϵ*Δ*u* + ϵΔ*uP*(*u*). The chance of no epidemic occurring up to time *t* is accordingly

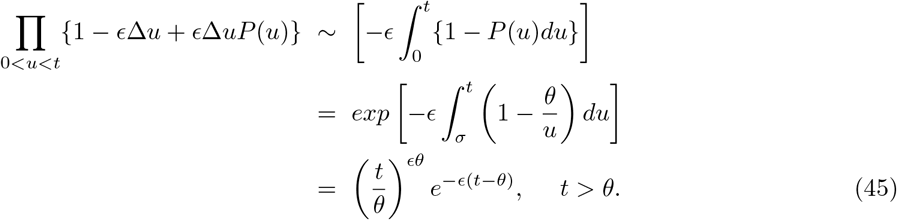

using (44). The lower limit of the integral in the last but one line above is *θ*, since *P*(*u*) = 1 when 0 *< u < θ* by virtue of (44). If we now write *F*(*t*) for the distribution function of time elapsing before the occurrence of a major epidemic, the quantity in (45) is precisely 1 − *g*(*t*). The corresponding frequency function, given by differentiating with respect to *t*, can therefore be written as

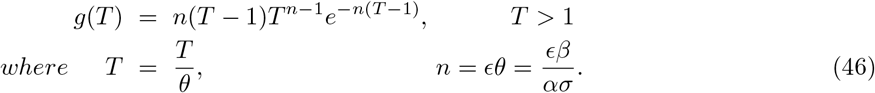

This distribution has a mode at *T*_*u*_ = 1 + *n*^−1*/*2^, and we might reasonably suppose that the mean 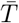 was approximately the same. It can be shown that 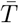 is relatively independent of *n* unless *n* is less than about 2. This means that the average renewal time 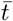 for major epidemics like Covid-19 is proportional to *θ*, but comparatively insensitive to changes in *ϵ* if the later is not too small.

## V. DISCUSSION

We shall now briefly discuss the important consequences of above mathematical expressions for Covid-19. Suppose we consider a relatively small community, in which new Covid-19 susceptibles either appear by actual birth or by introduction from outside. After a major outbreak of disease both the numbers of susceptibles and infectives will be low. At this point *ρ/b* is likely to be large, and extinction of infectives therefore highly probable. The population of Covid-19 susceptibles will continue to increase, and will reach a level where a new outbreak will easily occur if there are chance contacts with other infectives outside the community, or if these are allowed to join the group. We shall thus have a series of recurrent outbreaks of the disease. This behavior has been observed with the ongoing Covid-19 pandemic [3].

## VI. CONCLUSION

We have given a general mathematical treatment of Covid-19 as continuous time stochastic process. The chance of extinction and the consequences of introducing new Covid-19 infectives into the population was evaluated in general terms by using certain approximate arguments. We showed that the stochastic formulation of a recurrent epidemic like Covid-19 leads to the prediction of a permanent succession of undamped outbreaks of the disease. Finally, it must be mentioned here that in order to simplify the analysis we have omitted the consideration of the members of the community who are immune to Covid-19. A rigorous mathematical analysis must take this feature into account as well. However, we were able to derive certain useful conclusion without consideration of Covid-19 immune individuals. Other useful consideration would be to seek a way of influencing the rates of transitions between various classes of individuals. We have, however, omitted this aspect here. It should therefore be clear, at least in general terms, how the stochastic formulation of the model for a recurrent epidemic like Covid-19 leads to a permanent succession of undamped outbreaks of disease. It is hoped that the above analysis proves useful in the formulation of novel stochastic approaches to the ongoing pandemic of Covid-19.

## Notes

### Competing Interest Statement

The authors have declared no competing interest.

